# Biophysical Characterization of ParBS Condensates suggests a physical mechanism for segregation

**DOI:** 10.64898/2026.07.09.737391

**Authors:** Ritika Gupta, W. Seth Childers, Suleyman Ucuncuoglu, David Dunlap, Laura Finzi

## Abstract

The ParABS system orchestrates chromosome segregation in many bacterial species. Centromere-like parS sites nucleate binding of ParB, which subsequently spreads to adjacent non-specific DNA in the presence of CTP. Recruitment of additional ParB molecules and DNA looping via ParB-ParB interactions bring distal chromosome regions into proximity, while interaction with the ParA-ATPase motor drives segregation. In some bacterial species, ParB-parS complexes undergo phase separation into condensates, but their physico-chemical and mechanical properties remain poorly characterized. Turbidity measurements in the presence of CTP, DNA, and physiological mono and divalent salts showed that Mg2+ facilitates condensate formation, whereas K+ concentrations above ∼20 mM disfavor it. Microrheology measurements showed that ParB-parS DNA condensates are viscoelastic, with a room-temperature viscosity of ∼5 Pa·s and a network relaxation time of 0.1 s, allowing rapid responses to deformation. Fluorescence combined with force spectroscopy showed that mechanical disruption of ParB-DNA condensates requires 5–7.5 pN of DNA tension, lower than the force required to stall RNA polymerase but higher than that required for chromosome and plasmid relocation during segregation. These results suggest that ParB-parS condensates dynamically rearrange while maintaining sufficient cohesion to sustain segregation forces without interfering with genomic transactions.

## Introduction

Condensates formed by liquid-liquid phase separation enable cellular compartmentalization [1]–[4] without the high energetic cost of a membrane enclosure. They exhibit a wide range of physical states, from fluid and viscoelastic to more solid [5]–[7]. A fluid condensate allows rapid molecular exchange, whereas a rigid one promotes long-lived molecular interactions that may interfere with or arrest biochemical activities. Such different physical states have been implicated in biological processes like gene activation or repression [8], signaling modulation [3], and, in some cases, pathological protein aggregation [9], [10].

Since prokaryotes lack internal membrane-bound organelles, they rely on condensates to spatially and temporally coordinate key biochemical activities [4], [11]–[14]. In bacteria, condensates represent a quickly tunable system for cellular regulation [3], [4], [15]–[18].

Here, we focus on the *C. crescentus* ParABS chromosome segregation machinery, a condensate-forming system responsible for segregating newly replicated chromosomes and extra-chromosomal plasmids in many bacterial species [19]–[22]. The system is composed of the centromere-like parS DNA sites, the ParB-CTPase and the ParA-ATPase [23], [24]. CTP binding [25]–[29], facilitates ParB spreading beyond parS sites for approximately 10 kbp by driving the interaction between the NTD domains in the ParB dimer and inducing the transition to a clamp conformation which weakens the affinity of the dimer for the parS site. Subsequent CTP hydrolysis causes the opening of the clamp and facilitates ParB-ParB interactions in trans which leads to bridging of distant sites on the DNA [30]. The multivalent interaction between ParB and non-specific DNA leads to liquid-like nucleoprotein condensates [27], which can fuse on a time scale of 5±3 s [20].

The DNA-bound ParB is also known to interact with ParA. ParB stimulates the ATPase activity of ParA, which, in turn, causes the dissociation of ParA and the biased diffusion of the DNA-ParB nucleoprotein complex towards higher concentrations of ParA-ATP. This chemical force drives the directional motion of the newly replicated chromosome, and extrachromosomal plasmids, towards the pole of the cell [31]. The consequent drag force acts on a ParB-DNA condensate over a scale of several minutes which might disrupt the condensate. However, ParB-DNA foci within cells remain intact during movement suggesting that condensates retain structural integrity during segregation. This begs questions about (i) the parameters that favor condensate formation, (ii) how quickly the ParB-DNA condensates can reorganize in response to internal perturbations, such as those experienced during movement, and (iii) the chemical and mechanical stability of condensates when subject to a force load. To address the first question, turbidity measurements in different conditions of ionic strength and DNA concentration were used as proxies for phase separation and confirmed with imaging using fluorescently labeled ParB. While details of the ParB-DNA binding interaction, including the effect of permissive variants, have been recently characterized [29], the contributions of electrostatic interactions are incompletely characterized although their effects on phase separation are well-recognized [32], [33], [34]. Similarly, the dependence of ParB-DNA binding on DNA concentration needs further definition. To address this second question we considered condensates on several time scales: the sub-second timeframe of individual ParB proteins associating with parS sites through multivalent protein-DNA interactions, the seconds-to - minutes timeframe of the spreading of the nucleoprotein complex via cooperative ParB-ParB interactions [20], [31], and the bacterial chromosome segregation that spans tens of minutes, comparable to the cell lifetime. Thus, we measured the viscoelastic properties of condensates formed by ParB and a 6kbp DNA fragment from the C. *crescentus* genome containing 7 ParS sites (ParB-parS condensates) in order to define the network relaxation time reflecting rearrangements within condensates. A short relaxation time indicates a material that readily flows under stress, due to an ensemble of dynamic interactions, while a longer one indicates resistance to deformation, due to strong, long-lived molecular interactions. Therefore, the network relaxation time directly influences condensate mobility and nucleoprotein functions.

Additionally, as ParB-ParA interactions translate segments of DNA within ParB-DNA condensates, tensions of ∼4 fN [35] develop that could disrupt the condensates. Measuring the mechanical stability of condensates force is, therefore, relevant to understanding how they operate in chromosome segregation.

Here, we characterized the phase behavior, physical properties, and force-generating capacity of ParB-parS DNA condensates. We quantified their viscoelastic profile and estimated network relaxation timescales using passive microrheology in an optical tweezer setup. The tweezers were also used to measure the forces exerted by ParB on non-specific DNA. The results obtained provide a mechanical and physical framework for understanding how bacterial phase-separated condensates support essential genomic processes.

## Materials and Methods

### DNA constructs for turbidity and microrheology measurements

ParS-containing DNA – We used parS6000 which is a 5765 bp linear DNA containing seven parS sites. It was prepared by PCR using *Caulobacter crescentus* NA1000 genome as the template with primers otXYS13 and otXYS14 (sequences listed in Table S1) and KOD Hot Start DNA polymerase (MilliporeSigma Novagen), followed by gel purification. From now on, it will be referred to as parS DNA.

Non-specific DNA for turbidity measurements was prepared from a 5848 bp DNA plasmid (pDM N1 DNA) [36], which was linearized using the restriction enzyme BamHI from NEB according to the manufacturer’s protocol.

### Protein Purification

#### Expression and purification of His-ParB-mCherry-His

Plasmid pXIY0010 was transformed into *E. coli* DE3 Rosetta cells and plated on 100 µg/mL ampicillin and 60 µg/mL chloramphenicol LB agar and grown overnight at 37 °C. An overnight culture was produced from a single colony and then made into a freezer stock. A culture from the freezer stock was started and incubated at 37 °C overnight with LB media with 50 µg/mL ampicillin and 20 µg/mL chloramphenicol. The next morning, 22.5 mL of the overnight culture was inoculated to every 1 L LB media with appropriate antibiotics (12 L media in total). When the OD600 reached 0.4, the temperature of the incubator was set to 5 °C. When the real temperature of the incubator was lower than 30 °C, the expression of the protein was induced with 0.5 mM IPTG, then incubated at 30 °C for 3 hours. Cultures were centrifuged (4 °C, 4000×g, 10 min). Most of the supernatant was discarded keeping only around 5 mL, which was used to resuspend the cell pellets, then stored at -80 °C.

To purify, the resuspended cell pellets were thawed at 4 °C and resuspended in less than 200 mL lysis buffer (100 mM Tris-HCl, pH 8.0, 300 mM NaCl, 20 mM imidazole, 5% glycerol, 250 U universal nuclease, 5 ug/mL lysozyme, supplemented with a SIGMAFAST protease inhibitor tablet). Cells were passed through Emulsiflex C3 (Avestin) (15,000 psi, 50 min) before centrifugation (4 °C, 20,000×g, 35 min) to remove cell debris. The supernatant was loaded onto HisTrap FF column (GE Healthcare) and was eluted with elution buffer (200 mM imidazole). A260/A280 was checked by NanoDrop to pick the fractions whose percentage of nucleic acid was lower than 1%. The fractions were loaded onto Cytiva HiPrep Sephacryl S200 HR Size Exclusion Column 16/60 and eluted with dialysis buffer (50 mM Tris-HCl, pH 8.0, 100 mM NaCl, 5% glycerol), followed by ultra-centrifuge (4 °C, 70,000 rpm, 1 h). Fractions containing ParB-mCherry were concentrated using 30,000 MWCO Vivaspin centrifugal filters to 9.67 mg/mL, aliquoted, and stored at - 80 °C.

#### Expression and purification of GA-ParB

All the buffers are the same as for His-ParB-mCherry. GA-ParB-His was expressed from an *E. coli* strain which contains the plasmid pXYS031, and cell pellets were passed and centrifuged as for ParB-mCherry. After that, the supernatant was incubated with 200 µL Ni-NTA agarose resin slurry (ThermoFisher) per liter of culture with gentle rocking at 4 °C for 1 hour. The resin was separated by using a gravity-flow column, followed by washing with wash buffer and then B1 buffer (1 M NaCl, 50 mM Tris-HCl, pH 8). Then GA-ParB-His was eluted with elution buffer. A260/A280 was checked by NanoDrop to pick the fractions whose percentage of nucleic acid was lower than 1%. The fractions were concentrated with 30,000 MWCO Amicon centrifugal filters. To cleave the His-tag, 2 mg home-purified TEV protease was used.

### Biotinylation of λ-DNA

Bacteriophage λ DNA was labeled using the Klenow fragment according to the following protocol. A 50 μL reaction was assembled containing: 32.53 μL of 500 μg/ml λ DNA (New England Biolabs, Ipswich, MA), 5.05 μL nuclease-free water, 5 μL NEBuffer 2 (New England Biolabs), 0.19 μL dATP (10 mM), 0.19 μL dCTP (10 mM), 0.19 μL dGTP (10 mM), 1.85 μL Biotin-16-dUTP (AAT Bioquest, Pleasanton, CA) and 5 μL DNA Polymerase I, Large (Klenow) Fragment (New England Biolabs).

Reactions were incubated at 37 °C for 15 minutes with DNA Polymerase I, Large (Klenow) Fragment. The enzyme was inactivated by heating at 75 °C for 20 minutes, and samples were cooled to 10 °C. Labeled λ DNA was purified using the Zymo-Spin IC-XL column (Zymo Research, C1002-50) and DNA Clean & Concentrator kit (Zymo Research, D4013) according to the manufacturer’s instructions.

### Phase Separation and turbidity measurements

To assess the phase separation propensity of a ParB-parS DNA mixture, ParB was added to 20 µM final concentration in 50 mM Tris HCl (pH 7.5), 0, 5 or 10 mM MgCl_2_, 100 mM monopotassium glutamate (MPG), 2 mM CTP, and 8.5% (w/v) 8 kDa polyethylene glycol (PEG 8000) and varied ParS DNA concentrations of 1, 5, and 10 ng/µl. The solution containing protein and DNA was gently mixed by pipetting, and the turbidity was measured at 350 nm using a NanoDrop spectrophotometer (Thermo Fisher Scientific). Three technical replicates were prepared for each condition, and, in each case, fluorescence images were collected at multiple locations (Figure S1). Intensity profiles were obtained using the plot profile function of Fiji[37](Figure S2). The parS concentration corresponding to the peak value of the plot of turbidity versus parS concentration was chosen to further examine the effect of MPG.

### Microrheology measurements

ParB-parS DNA condensates were prepared at 40 µM ParB and 10 ng/µl parS DNA in a buffer containing 50 mM Tris HCl (pH 7.5), 10 mM MgCl_2_, 25 mM MPG, 2 mM CTP, 8.5% (w/v) PEG 8000, and carboxylate beads of 1 μm in diameter (Spherotech, Lake Forest, IL). The sample was deposited on a BSA-coated coverslip and covered with a glass slide using two layers of double-sided tape. To avoid evaporation, the open ends were sealed with mineral oil. After ∼ 2 hours, condensate droplets settled onto the coverslip. Many droplets contained one bead and had a minimum size ∼10 µm. The beads were trapped using an IR laser of wavelength 1044 nm in the *z* plane. The laser power was adjusted to only trap the bead and not the condensate.

Eight beads were trapped in different condensates across multiple samples, and their motion was tracked for ∼20-30 mins in the *xy* plane. The bead trajectory was analyzed independently in the x and y planes to determine the trap stiffnesses, *k*_x_ and *k*_y,_ respectively.

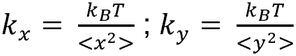

Where *k_B_* is the Boltzmann constant and *T* is the temperature in degrees Kelvin. Following the procedure described in [38], [39], the normalized position autocorrelation function A(ω), where ω is the frequency, was calculated and, subsequently, the real and imaginary rheological moduli were determined as:

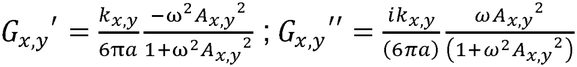

*G*’ and *G*” correspond to the storage and loss modulus, respectively and a is the radius of the carboxylate bead used. For each bead, *G*’ and *G*” in *x* and *y* were calculated and averaged over frequency. Since the data were collected using a bright-field camera at 80 Hz, the reported complex moduli are shown up to 40 Hz to avoid artifacts arising from the Nyquist limit [40]. The frequency-dependent viscosity at *T*_*room*_ was calculated using

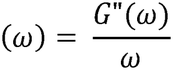

Autocorrelation data and viscoelastic moduli of each bead, and more details about the data analysis are shown in Figures S3 and 4.

### Single-molecule measurements

Single-molecule experiments were conducted using a C-trap microscope (Lumicks, Amsterdam, Netherlands) equipped with a uFlux microfluidics flow chamber. Two 4.35 μm streptavidin-coated beads (Spherotech) were captured in the bead channel, calibrated, and transferred to the DNA channel (20 mM Tris HCl, 50 mM MPG, 4 mM MgCl_2_, as described in [41]), containing 10 pM λ-DNA. To establish a tether, one bead was held stationary, while the other was moved back and forth until a distinct increase in force was observed upon separation. Then, the beads were immediately brought closer together, the flow was stopped, and the bead-DNA-bead assembly was transferred to the buffer channel (20 mM Tris HCl, 50 mM MPG, 4 mM MgCl_2_). Repeated stretching-and-relaxation cycles were performed to measure force as a function of distance between beads (F-D curves). The presence of a single DNA tether between the beads was confirmed by fitting the data to the Worm-like Chain (WLC) model (Figure S5). In the buffer channel, the tethered DNA was stretched with force up to ∼14-18 pN before moving it to the protein channel (20 mM Tris HCl, 50 mM MPG, 4 mM MgCl_2_, 0, 0.5 or 2mM CTP, 200 nM ParB, and 8% PEG8000 as indicated). The stretching was to prevent excessive protein-induced condensation upon entry in the protein channel. The DNA was repeatedly stretched and relaxed to record F-D curves and detect condensates. Three different DNA molecules were examined in this way for each experimental condition (Figure S6). For fluorescence measurements, the flow, perpendicular to the DNA tether, was turned ON to a minimal value of 0.1 bar, but the F-D curves were recorded without any flow.

## Results

### Phase Behavior of ParB Depends on parS DNA and Salt Concentration

The packaging by proteins of genomic DNA in the bacterial cells is a dynamic process that directly governs DNA access for vital processes such as transcription, repair, etc. Increasing evidence indicates that condensates also play a role in packaging and genome processing, suggesting that their assembly and disassembly must be precisely controlled [3], [42]. Therefore, identifying the determinants of condensation and de-condensation is critical. Thus, we investigated condensate formation as a function of DNA and monovalent, as well as divalent, ion concentration. To do so, we utilized solution turbidity at 350 nm as a readout for phase separation, a ∼6kb linear DNA fragment (see Materials and Methods) containing 7 parS-like sites (parS DNA), mCherry-tagged ParB protein and PEG8000 as a crowder to mimic the *in vivo* environment. We chose to vary K^+^ (monopotassium glutamate, MPG) and Mg^2+^ (MgCl□), because they are physiologically relevant intracellular cations.

We fixed the concentration of ParB, CTP and MPG at 20 µM, 2 mM and 100 mM respectively, and varied the concentration of parS DNA from 1 to 10 ng/µL, while maintaining the buffer and crowder conditions constant. We repeated such parS DNA titration in the presence of 5 and 10 mM Mg^2+^. Low turbidity was observed across all parS DNA concentrations in the absence, or in the presence of 5 mM MgCl_2_ (Figure 1a) indicating the presence of few small condensates. However, in the presence of 10 mM MgCl□, turbidity initially increased, peaked, and then decreased as the concentration of parS DNA was increased (Figure 1a). In contrast, increasing concentrations of MPG from 25 mM to 300 mM while maintaining parS DNA at 5 ng/µl and Mg^2+^ at 10 mM suppressed phase separation (Figure 1b). These latter observations on the effect of K^+^ are in-line with past studies showing that NaCl attenuated *Caulobacter crescentus* ParB phase separation [43]. The distinct dependencies on Mg^2+^ and K^+^ concentrations highlight complex salt-specific modulation of phase separations of ParB-parS DNA. The concentrations of K^+^ within the bacterial cytoplasm is ∼200 mM [44], which suggests that ParB-parS DNA-mediated condensation in the cellular context critically depends on Mg^2+^ compensating the negative effect of K^+^. Since physiological concentrations of free Mg^2+^ in *E. coli*, range between 1 and 10 mM [45], this may indeed be the case.

**Figure 1.**
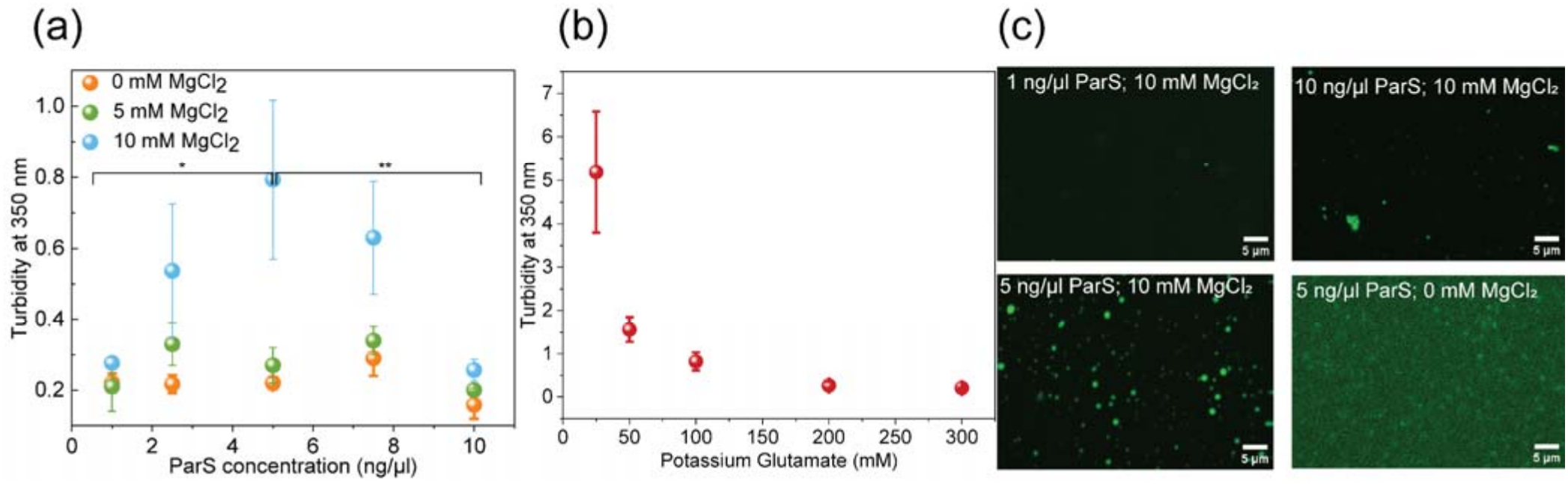
The dependence of ParB-parS DNA condensates on parS-DNA and salt concentrations. (a) Turbidity values at 350 nm of ParB-parS-DNA solutions with varying parS-DNA concentration in the presence of 20 µM ParB, 2 mM CTP, 100 mM MPG and 0, 5 and 10 mM MgCl_2_. (b) Solution turbidity at fixed ParB (20 µM)) and parS DNA concentration (5 ng/µl) and varying monopotassium glutamate (MPG) from 25 to 300mM. Statistically significant differences calculated by One-Way ANOVA + Tukey’s HSD: * p<0.05, ** p<0.01. (c) Representative confocal fluorescence images of the ParB-parS DNA mixture in the presence of 20 µM ParB and various conditions of salt and parS-DNA.

For visual confirmation, we used confocal fluorescence microscopy of mixtures as described in the turbidity assays. Condensates were observed in the same conditions that produced trubidity, and none were detected under conditions that were unfavorable to phase separation (Figure 1c). these results suggest that Mg^2+^ ions screen electrostatic repulsions between DNA segments, allowing them to condense.

### ParB-parS DNA condensates are viscoelastic

Figure 2a schematically illustrates the key components of the ParB-parS DNA condensates used in the study: at the top, the centromere-like parS site highlighted in yellow and at the bottom the ParB main components, namely, an N-terminal and a CTP binding domain, a Helix-turn-helix (HTH) DNA binding domain and a C-terminal dimerization domain. Having established the conditions of parS DNA and salt concentrations that yield the highest degree of turbidity at 350 nm (Figure 1), we investigated the time-dependent material properties of the condensates under these phase separating conditions. After embedding 1 μm carboxylate beads inside the ParB-parS DNA condensates (average diameter ≥ 10 μm), we performed passive microrheology measurements. It was important to ascertain that ParB did not adsorb on the carboxylate beads inside condensates and enrich ParB in the surrounding environment (Figure S7). Then, one bead per condensate was trapped using the IR laser (1064 nm) of the C-trap microscope (Figure 2b, left) and its motion in the *xy* plane was tracked for 15-30 mins. The different colors in the map in the right panel of Figure 2b represent the trajectory of a representative bead at different time intervals during the tracking. The random spatial distribution of such trajectories confirms random diffusion of the bead and material homogeneity of the condensate. The data also reveal that the viscoelastic moduli of ParB-parS DNA condensates are dependent on the frequency of the perturbation (Figure 2c). The storage modulus, *G*’, which represents the elastic portion of the viscoelastic behavior, describes the solid-state behavior of the condensate. The loss modulus *G*’’ which characterizes the viscous portion of the viscoelastic behavior, describes the liquid-state behavior of the sample. At low frequencies, or over long timescales (above 25 s), the loss modulus dominates the storage modulus, indicating viscous, or liquid-like, behavior. At 0.1 s, the values of *G*’ and *G*’’ become equal indicating that the condensate displays both solid- and liquid-like behavior. Note that ParB condensates take ∼5 seconds to fuse as seen in *E. coli* cells [20]. This implies that re-equilibration after perturbation must involve molecular rearrangements that occur faster than those involved in condensate fusion.

**Figure 2.**
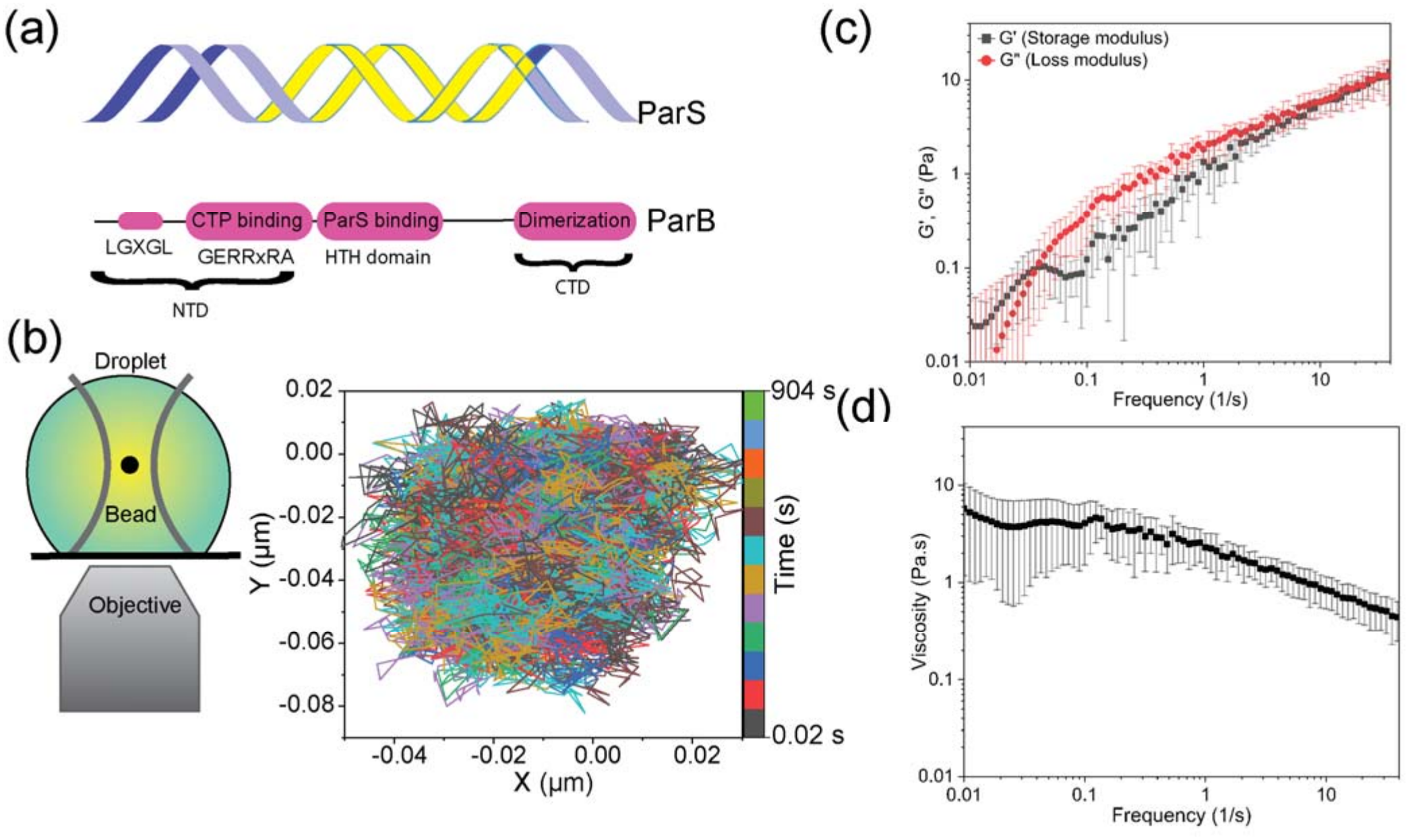
ParB-parS condensates are viscoelastic. (a) Schematic of a 16 bp ParS site (highlighted in yellow) and the domain architecture of ParB. (b) Schematic of passive microrheology measurements using optical tweezers and the representative motion trajectory of a trapped bead in xy plane. (c) Averaged frequency-dependent storage and loss moduli of ParB-parS condensates. (d) Frequency-dependent viscosity of ParB-parS condensates extracted from bead tracking.

Dividing the loss modulus by the frequency, we calculated frequency-dependent viscosities. The low-frequency viscosity, ∼5 Pa·s, is comparable to the zero-shear viscosity, used as an indicator of the stability of the condensate (Figure 2d). This value is comparable to that of other condensates with similar network relaxation time [46].

### ParB generates force on DNA

After characterizing the viscoelasticity of ParB-parS DNA condensates, we sought to visualize the condensates and characterize their dependence on DNA tension. For this measurement, λ-DNA was tethered between two optically trapped beads (Figure 3a) and the tension was monitored during stretching and relaxation in the presence of ParB with and without CTP (Figure 3b and SI movies 1 & 2). ParB is known to bind non-specifically to λ DNA and form condensates [47], and we assume that their physical characteristics are similar to those of the ParB-parS-DNA condensates. ParB-λ DNA condensates were force dependent in the absence of CTP, did not form in the presence of 0.5 mM CTP, while they formed and resisted tension in the presence of 2 mM CTP.

**Figure 3.**
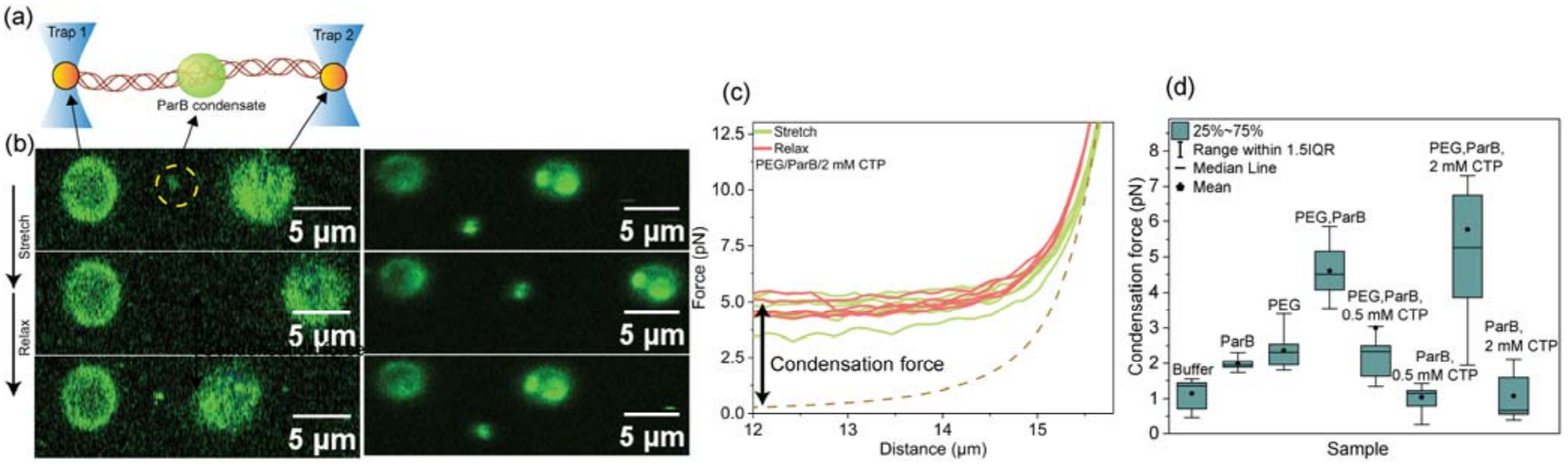
Visualization and quantification of the force exerted by ParB condensates on λ DNA. (a) Schematic illustration of the dual-trap optical tweezers setup. A single λ DNA molecule is tethered between two trapped beads, and ParB molecules condense the DNA (green). (b) Representative fluorescence snapshots from SI movies 1 and 2 showing the PEG/ParB condensate during a stretch-relax cycle in the absence of CTP (Left) and in the presence of 2 mM CTP (Right). Scale bar = 5 µm. All videos were recorded during orthogonal flow. (c) Representative force-distance (F-D) curves obtained during the stretching (green) and relaxation (red) cycle in the presence of PEG+ParB. The dashed brown line represents the theoretical force-extension curve of naked DNA (worm-like chain model). The vertical arrow indicates the magnitude of the condensation force generated by the condensate. All F-D curves were recorded in the absence of flow. (d) Box plot comparing the condensation forces measured under different conditions. All measurements were conducted in the buffer indicated in Materials and Methods.

To quantify the contractile force on the DNA within the ParB-λ DNA condensates (condensation force), the DNA was stretched to approximately 16 μm in the buffer channel before transferring it to the ParB-containing channel (see Materials and Methods). Through repeated cycles of relaxation and stretching from 12 to 16 μm in the ParB channel, the force at an extension of 12 μm was constant at 5-6 pN in the presence of PEG and 2mM CTP for three different tethers (Figures 3c,d and S6a). To distinguish the contribution of ParB from the crowder (PEG) and to better define the role of CTP in DNA condensation, force versus distance (F-D) curves were measured in buffer or in buffer supplemented with just ParB, or PEG, with and without CTP (Figures 3d and S6d,e). The presence of either PEG or ParB, individually and without CTP, compacted DNA with a force ∼2 pN. Instead, PEG and ParB together without CTP produced a condensation force around 5 pN (Figure S6c). When 0.5 mM CTP was added, the condensation force decreased to 2 pN (Figure S6e). In the absence of PEG, however, ParB with either 0.5 or 2 mM CTP generated only approximately 1 pN of condensation force, similar to ParB in buffer alone. Figure 3d summarizes the force values measured under these conditions when the beads in the C-Trap microscope were separated by 12 μm. These results are consistent with the dependence of condensate formation on CTP reported by Babl *et al*. [43] and with our own finding that 1–4 mM CTP of ParB-mCherry was necessary for robust phase separation in bulk (not shown).

## Discussion

### Phase separation is finely tuned by parS DNA, Mg^2+^ and MPG concentration

The formation of condensates of ParB and DNA shown here is consistent with previous evidence of phase separation of ParB and DNA, either containing specific ParB binding sites (parS) or not [20], [47]. However, this study extends the chemo-physical characterization of such condensation behavior focusing on the role of physiologically relevant cations (K^+^ and Mg^2+^), their valency and concentration. It further investigates the role of the concentration of parS DNA. The results indicate that the propensity of ParB-parS DNA to form condensates depends critically on salt valency and the concentration of both parS DNA and cations.

First, the data show that in the presence of 10 mM Mg^2+^, condensate formation is optimal at 5 ng/µl parS-DNA, which is the estimated concentration in the bacterial cell of the 6 kbp parS-containing chromosomal fragment we used, ∼6 ng/µl. Similarly, the 20 µM ParB used in the experiments is within the range of physiological concentrations in the cell [48]. In the absence of, or with 5 mM Mg^2+^, the turbidity was much lower (0.2-0.3, rather than 0.8), indicating fewer and smaller condensates. This level of condensate formation also occurred with non-specific λ DNA with similarly low Mg^2+^ in single-molecule C-Trap measurements.

While divalent Mg^2+^ ions, above 5 mM, strongly enhance phase separation, monovalent MPG concentration beyond 25 mM is inversely related to phase separation. The dependence of ParB-parS DNA condensate formation on MPG is similar to that reported for Na^+^ [43], although sodium ions are at a much lower in the bacterial cell than potassium ions. Given that Mg^2+^ and glutamate are the two most abundant divalent ions in bacterial cells, and that we used a concentration consistent with that of free Mg^2+^ in the cell, their effects on condensate formation support the idea that electrostatics play a central role in regulating the ParB-parS interaction. Unlike CTP, which functions as a conformational switch responsible for ParB binding and spreading to non-specific DNA, the well-known charge-screening and DNA compacting abilities of the Mg^2+^ cation are likely synergistic with the long-range DNA bridging capability of ParB in forming higher-order nucleoprotein complexes. This observation is in line with other reports suggesting Mg^2+^ favors condensate formation [49].

A previous study has shown that divalent ion variations continuously tune the microenvironments and fluid properties of heterotypic and homotypic RNA droplets and the authors speculated that those results indicate a general mechanism for modulating the biochemical environment of RNA coacervates in the cellular context [50]. Similarly, we assume that variations in Mg^2+^ may tune the ParB-parS DNA condensates depending on cellular requirements.

### A viscous nature and a fast relaxation time maintain integrity of ParB-parS DNA condensates

The viscoelastic properties of ParB-parS DNA condensates are consistent with Maxwell-type fluids, which are characterized by both (i) viscosity and (ii) the capacity to recover after a deformation, an elastic behavior more typical of solids than liquids. For the ParB-parS DNA condensates at T_room_, we measured a viscosity of 5 Pa·s in the absence of shear (low frequency), comparable to that of corn syrup in similar conditions. This very high viscosity suggests a dense network of molecular interactions. Such a network is consistent with the measured relaxation time of ∼0.1 s, which indicates that they dynamically remodel on a temporal scale consistent with that of molecular interactions corresponding to chromosome segregation. Since these condensates are due to specific and non-specific ParB-DNA interactions and the many DNA-bound ParB-ParB interactions and, fast recovery after deformation is likely due to the ability to quickly re-establish ParB-DNA complexes, as well as ParB-ParB-mediated DNA loops [30]. This idea is supported by the fact that ParB dimers remain bound to DNA for an average time of 76 ± 2 seconds (in *B. subtilis*), with CTP hydrolysis acting as the limiting factor, and once loaded on non-specific DNA in the presence of CTP, dimers are mobile, with diffusion coefficients around 0.1–0.4 µm^2^/s [41]. ParB bound to parS is, instead, almost immobile, 0.05 µm^2^/s [51]. Previous work shows that macroscopic fusion of condensates of ParB and parS DNA that form at different locations in the *E. coli* nucleoid occurs within ∼5 seconds [20]. Both the relaxation and fusion time scales of these condensates are orders of magnitude faster than the ∼20-40 minute timescale of chromosome segregation and cell division [52], [53], underscoring the importance of their dynamics in cell function. Given that the condensates, packaging the newly-replicated chromosome for segregation, must sustain the tension driving them to the cell pole without falling apart, their short relaxation time suggests that ParB-DNA condensates are poised to adapt to, and re-equilibrate after, any deformations due to their movement. Unlike solid aggregates, the liquid-like nature of ParB condensates in the crowded cell enables rapid molecular exchange, reversible CTP-dependent assembly and disassembly, and dynamic repositioning as replicated DNA molecules are segregated.

### ParBS condensates are pliable and robust

In bacterial chromosome segregation, ParB is a CTP-dependent DNA clamp that cycles through loading in the open conformation, spreading and condensation in the closed clamp conformation and opening after slow CTP hydrolysis to release DNA. Then, the cycle repeats. The ParB-CTP cycle regulates condensate formation from a biochemical point of view, as CTP binding switches ParB into a high-affinity multivalent state that promotes cooperative ParB– ParB interactions and liquid-like partition of parS DNA. CTP hydrolysis limits condensate size and maintains dynamic turnover. In this work, we studied the unexplored effect of tension on ParB-DNA condensates, as a function of CTP and PEG, investigating their mechanical regulation.

Comparison of force versus distance (F-D) curves of λ DNA recorded in buffer or in buffer supplemented with just ParB or PEG with and without CTP (Figures 3d and S6b,c) showed that crowding by PEG is necessary to obtain condensates with higher connectivity and highest condensation force. Additionally, Figure 3d shows that CTP concentrations sufficiently high to avoid depletion also produce ParB/PEG condensates characterized by high connectivity and exerting a high condensation force on DNA. These results are consistent with the dependence of condensate formation on CTP reported by Babl et al [43] and with our own finding that 1–4 mM CTP of ParB-mCherry was necessary for robust phase separation in bulk (not shown).

Notably, the ParB-λ DNA condensates in the absence of CTP, but presence of PEG, forms reversible liquid-like condensates on λ DNA, a process strictly modulated by the tension exerted on the DNA tether. Instead, no condensates were observed in the presence of 0.5 mM CTP, while 2 mM CTP favored and stabilized condensates against tension, impeding their disruption. This dependence of the stability of condensates against tension on CTP concentration might be explained considering the following.

Lambda only contains five parS pseudo sites each with four mismatches to a canonical TGTTTCACGTGAAACA sequence [54]. In the absence of CTP, ParB dimers bound to pseudo parS sites will not transition to the closed clamp conformation. Therefore, they will be less mobile and remain available for interactions with other ParB dimers bound on different DNA segments. Such interactions are weaker and easy to break when the DNA substrate is under tension. In the presence of 0.5 mM CTP, some ParB will bind CTP, form a closed clamp, diffuse and bridge distant DNA segments. CTP hydrolysis will then open the clamp and cause the dissociation of ParB with a low probability to bind CTP and repeat the cycle. CTP will be quickly depleted and condensates will not reform stably enough to be detected in our measurements. In the presence of 2 mM CTP, however, strong condensates may form with many inter-segmental interactions mediated by ParB proteins bound at several loci and spread over long distances. CTP is saturating, so that after hydrolysis, the ParB cycle can continue by binding a new CTP molecule.

Note that in bulk control measurements, a non-specific DNA fragment, which only has five parS pseudo sites with six mismatches to the canonical sequence, in buffer containing 4 mM Mg^2+^ and 2 mM CTP showed lower turbidity values throughout the range of DNA concentrations explored (Figure S8). However, at 5 ng/ul DNA, the turbidity equaled that of the same concentration of parS DNA in 5 mM Mg^2+^ and 2 mM CTP (Figure 1a).

Although the intracellular concentration of CTP in gram negative bacteria such as *E. coli* or *C. crescentus* is estimated in the sub-millimolar range, in average around 0.1-0.5 mM, its spatial distribution varies greatly [55]. The variability of the tension sustained by condensates in different conditions of CTP indicate small molecule regulation of susceptibility to forces in the cell and, in particular, to tensions acting on the chromosome and extrachromosomal plasmids. The tunability of resistance of the ParB-parS DNA condensates to force likely influences formation and dissolution according to different needs during the cell cycle, promoting segregation in the appropriate time window. For example, in mycobacteria, the bacterial chromosome replicates with a single ParB foci that divides to produce foci for each chromosome just prior to segregation [56]. The foci disappear shortly after segregation. Assuming that the foci correspond to ParB-DNA condensates, our F-D measurements suggest that to split or rupture the foci [57], tension on the DNA greater than 2.5-6 pN must be exerted on the foci depending on the CTP concentration. This range of forces is significantly lower than the stall force of motor enzymes such as RNA polymerase (∼25 pN) [58], suggesting that transcriptional machinery can draw DNA from the condensate, which therefore is not a significant steric roadblock to gene expression. This is in agreement with reported transcription elongation measurements through ParB condensates [59]. Indeed, the region flanking the parS site contains essential genes that must be transcribed. Thus, the liquid like properties of ParB-DNA condensates may have evolved to avoid impeding the transcriptional machinery.

Conversely, the few pN of tension sustained by a condensate is orders of magnitude higher than the estimated 4 fN drag forces involved in chromosome and plasmid segregation [35].

This ensures that condensates remain structurally intact under the mechanical loads required for partitioning and suggests the reason why a condensate may be the most efficient way to carry out segregation of the chromosomes. Just as an efficient way to move a garden hose is to grab it in different places and drag several loops, rather than pulling from one end, it may most effective to gather the bacterial chromosome into a condensate rather than stretching it out and recoiling it again at the pole (Figure 4). Such a model is compatible with currently accepted ones that focus on the molecular interactions of condensation [60], [61], and provides a mechanical mechanism for the translocation of the whole chromosome/plasmid.

**Figure 4.**
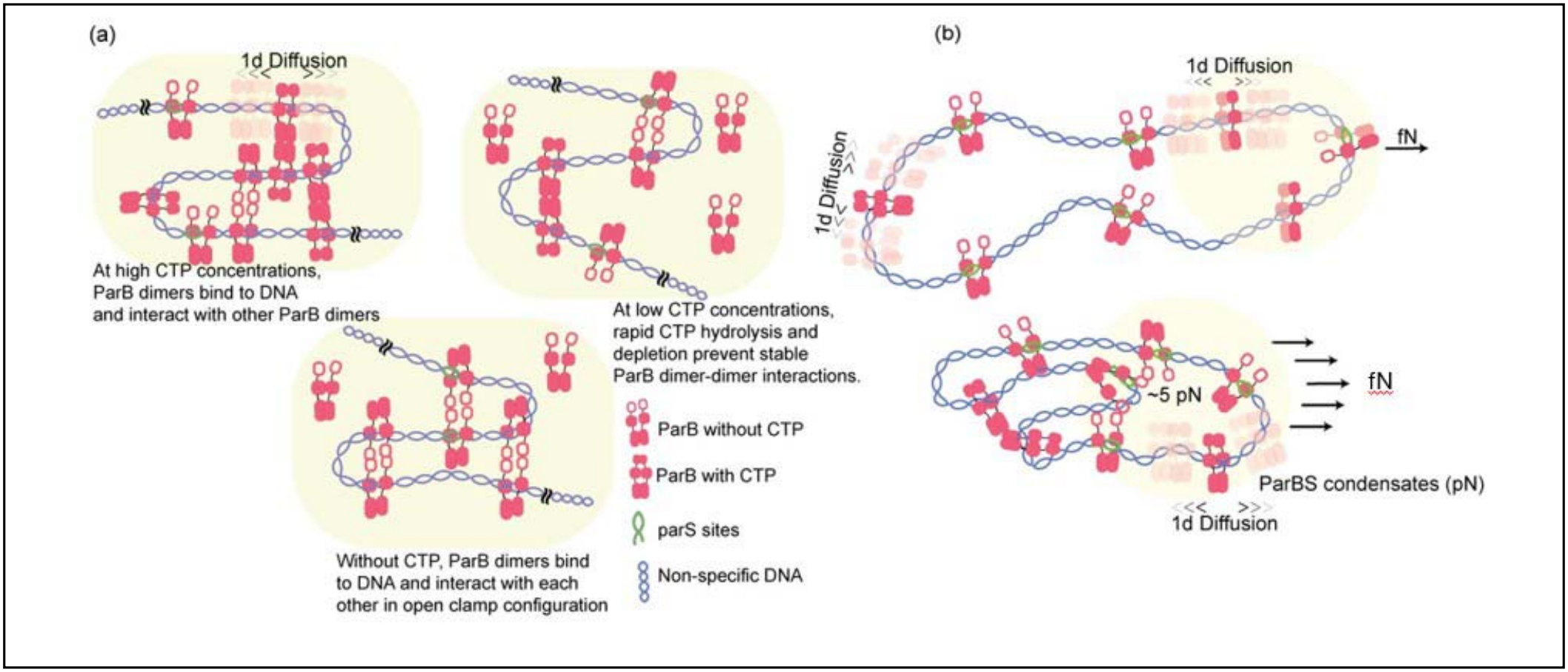
(a) Condensation is tuned by CTP and facilitates the segregation of the chromosome. (b) Schematic of garden hose mechanism for chromosome segregation.

Taken together, these results suggest that the ParB-DNA condensate system strikes a balance between the fluidity of liquids and the elasticity of solids, in order to successfully segregate genetic material. Shedding light on the role of ionic strength, molecular interactions, condensate material and mechanical properties, advances our understanding of the role of ParB condensates in the reorganization and relocation of the bacterial genome.

## Supplementary Data statement

Supplementary Data will be made available at NAR Online.

## Data availability

The data underlying the findings of this study will be made available on a public repository at publication.

## Author contributions

Ritika Gupta performed all experiments, generated figures and wrote the manuscript; Suleyman Ucuncuoglu labeled λ DNA with biotin; W. Seth Childers shared the ParB protein fluorescently labeled and not, parS6000 from Caulobacter crescentus, helped design the reseach and wrote the manuscript, David Dunlap helped design the research and wrote the manuscript, Laura Finzi designed and sponsored research and wrote the manuscript.

## Acknowledgments

We thank the Sanabria group for providing access to the Nanodrop spectrophotometer. This work was supported by NIH award 7R35GM149296 to LF, Clemson University startup to LF, and R01GM136863 from NIH NIGMS and the University of Pittsburgh Start-up funds to WSC. We thank Xinyue Song for purification of the ParB protein and parS.

