## Supplementary material for "Biophysical Characterization of ParBS Condensates suggests a physical mechanism for segregation": Figure S

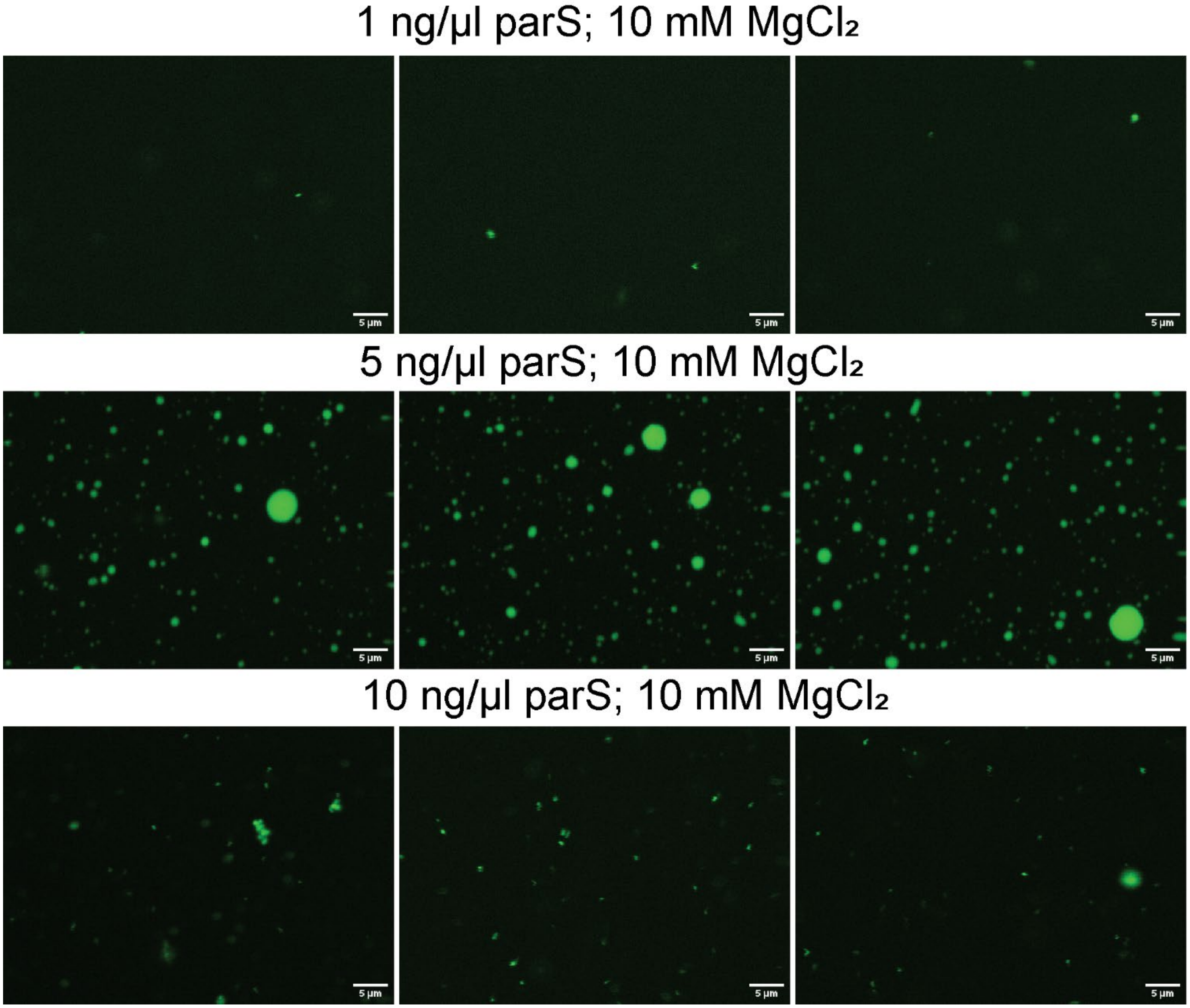


Figure S1. Three technical replicates of the fluorescence image of the ParB-parS DNA mixture (50 mM Tris HCl, 100 mM MPG, 2 mM CTP, and 8.5% PEG 8000) at three different concentrations of parS DNA and 10 mM $Mg^{2+}$.

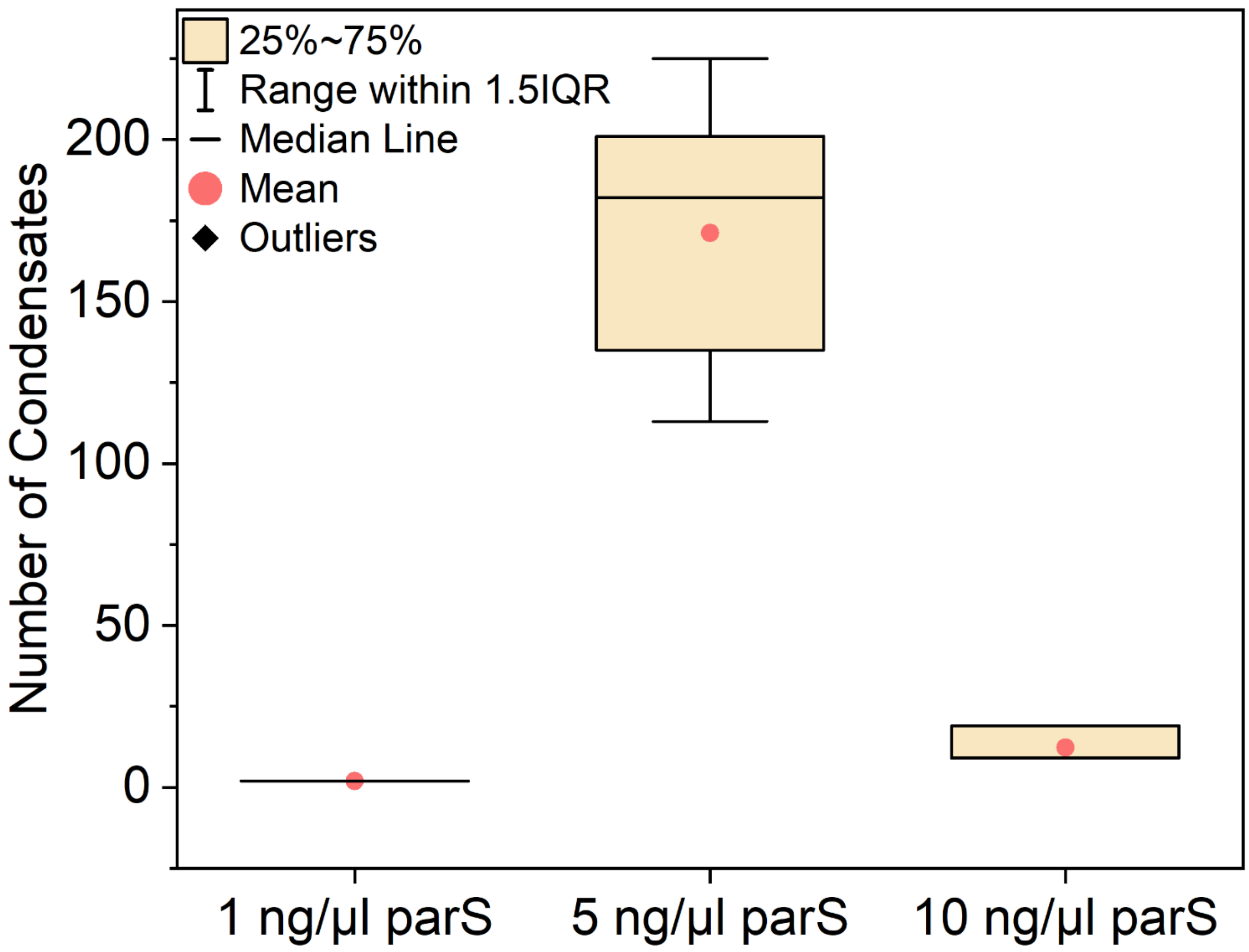


Figure S2. Quantification of the fluorescence/condensate level in the different experimental conditions indicated in Figure S1.

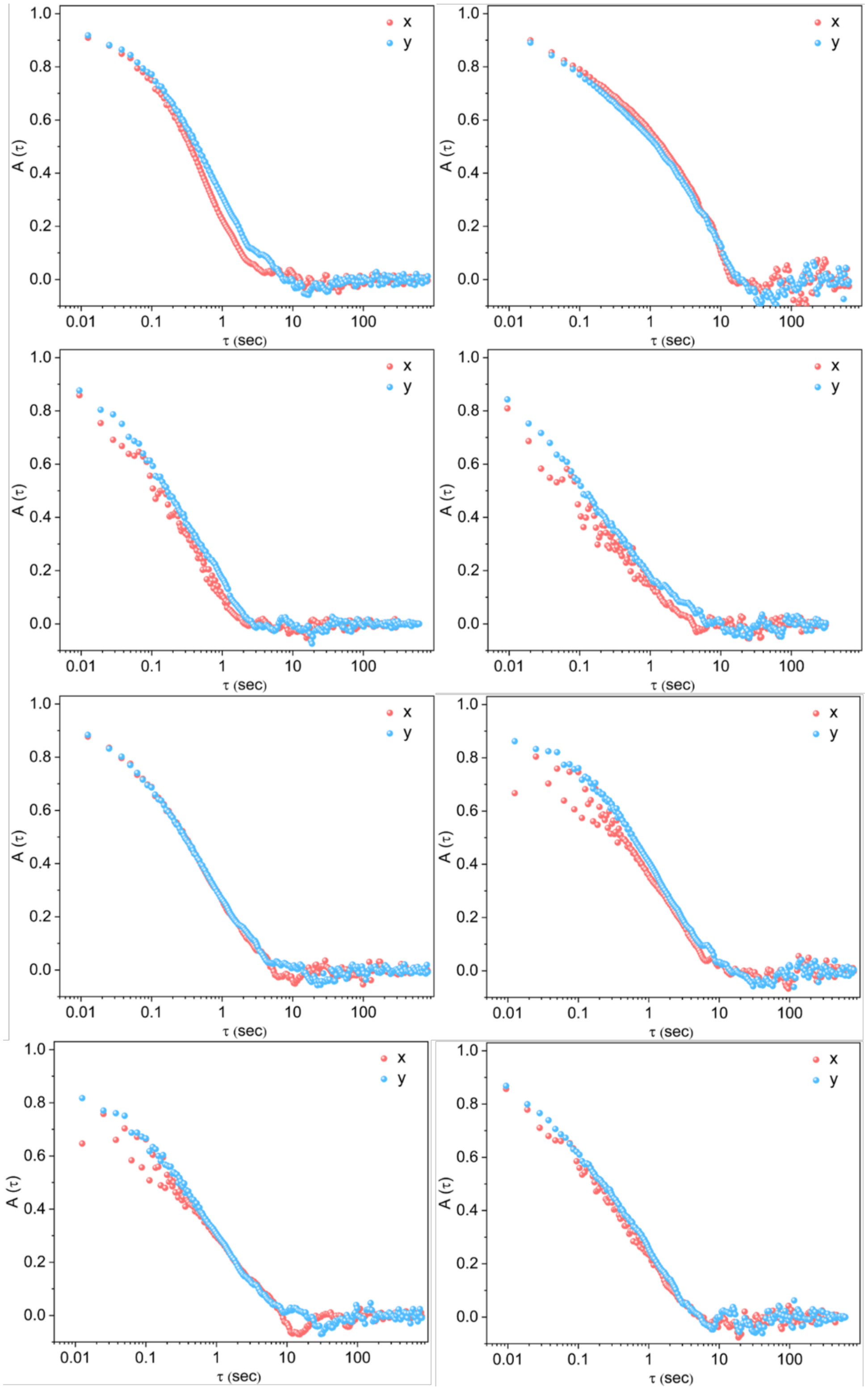


Figure S3. Calculated normalized position autocorrelation function vs lag time of different beads. Red and blue dots represent A(τ) of a trapped bead in x and y. The decay of A(τ) from 1 to 0 indicates the timescale over which the beads' motions become uncorrelated due to thermal diffusion. The tight overlap of the x and y traces demonstrates predominantly isotropic motion of the trapped bead.

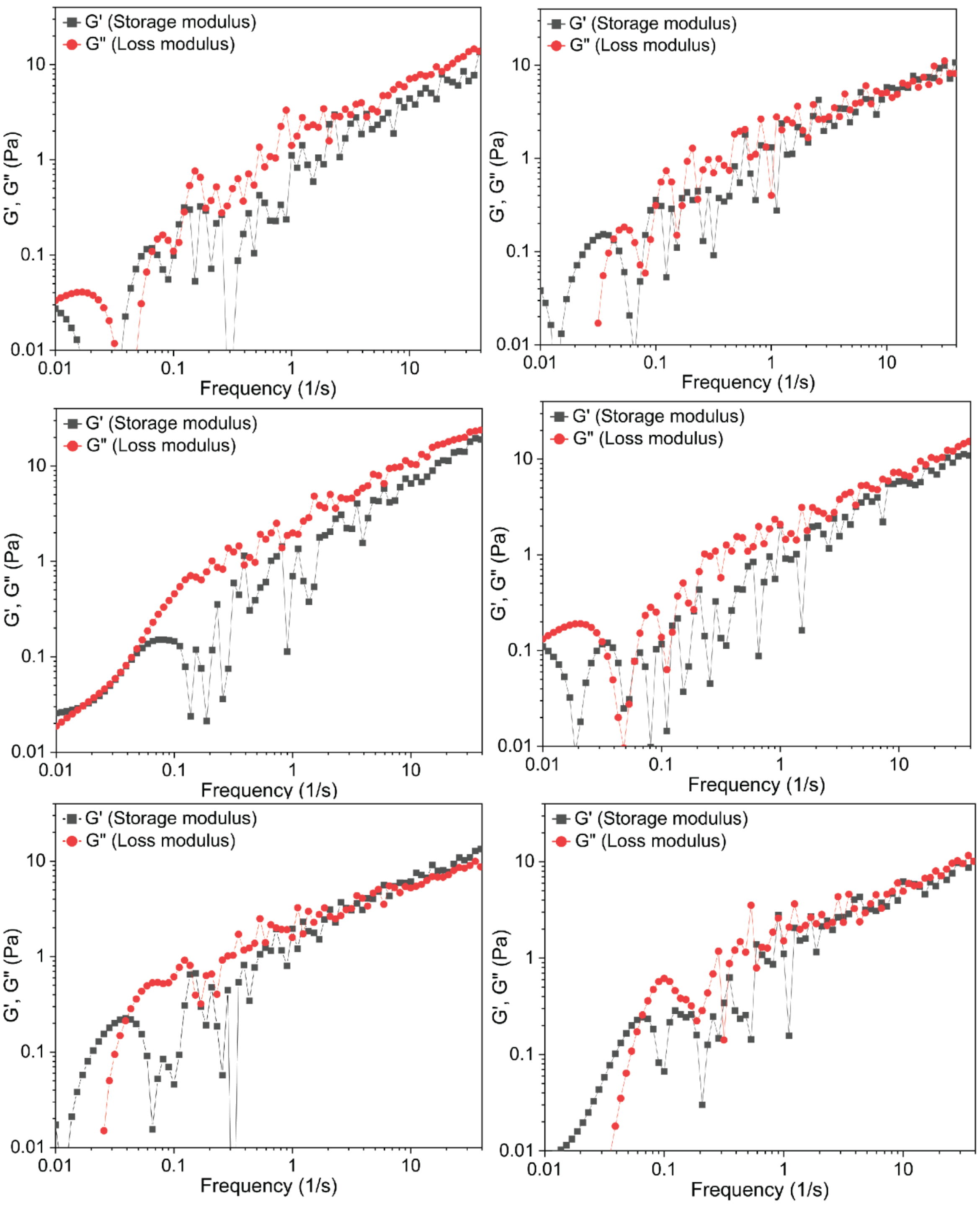


Figure S4. Variation of storage and loss modulus as a function of frequency in x and y for 3 representative beads. Across the probed frequency, the loss modulus remains higher than the storage modulus indicating the condensates exhibit liquid-like behavior.

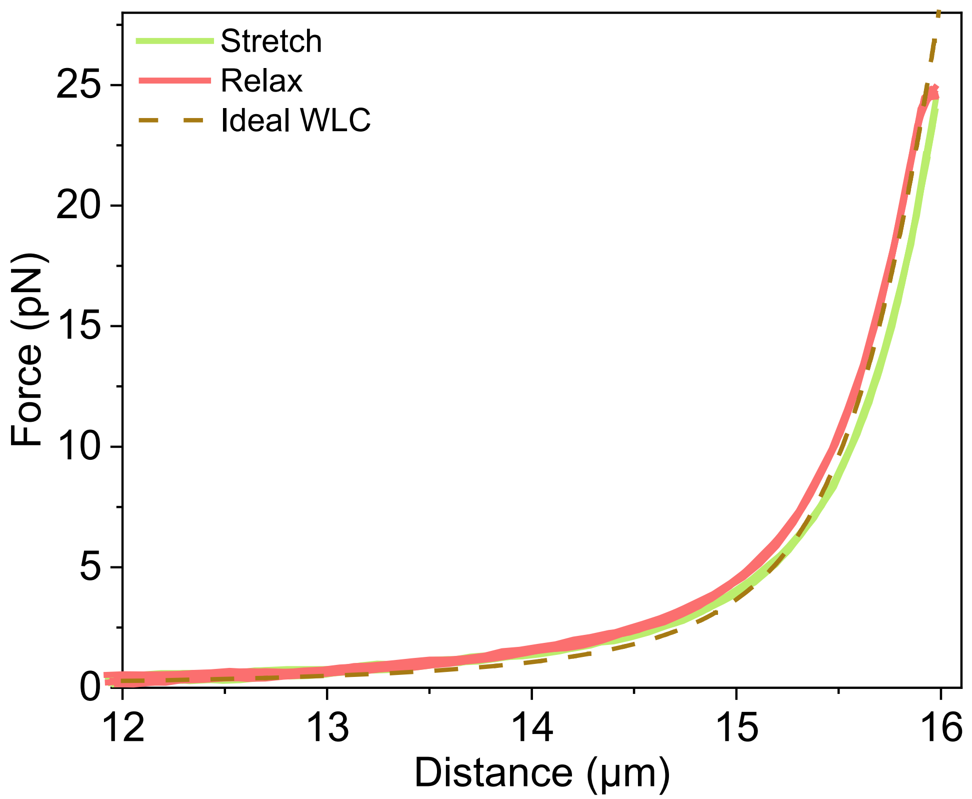


Figure S5. Representative FD curve of a DNA in the buffer channel confirming a single tether.

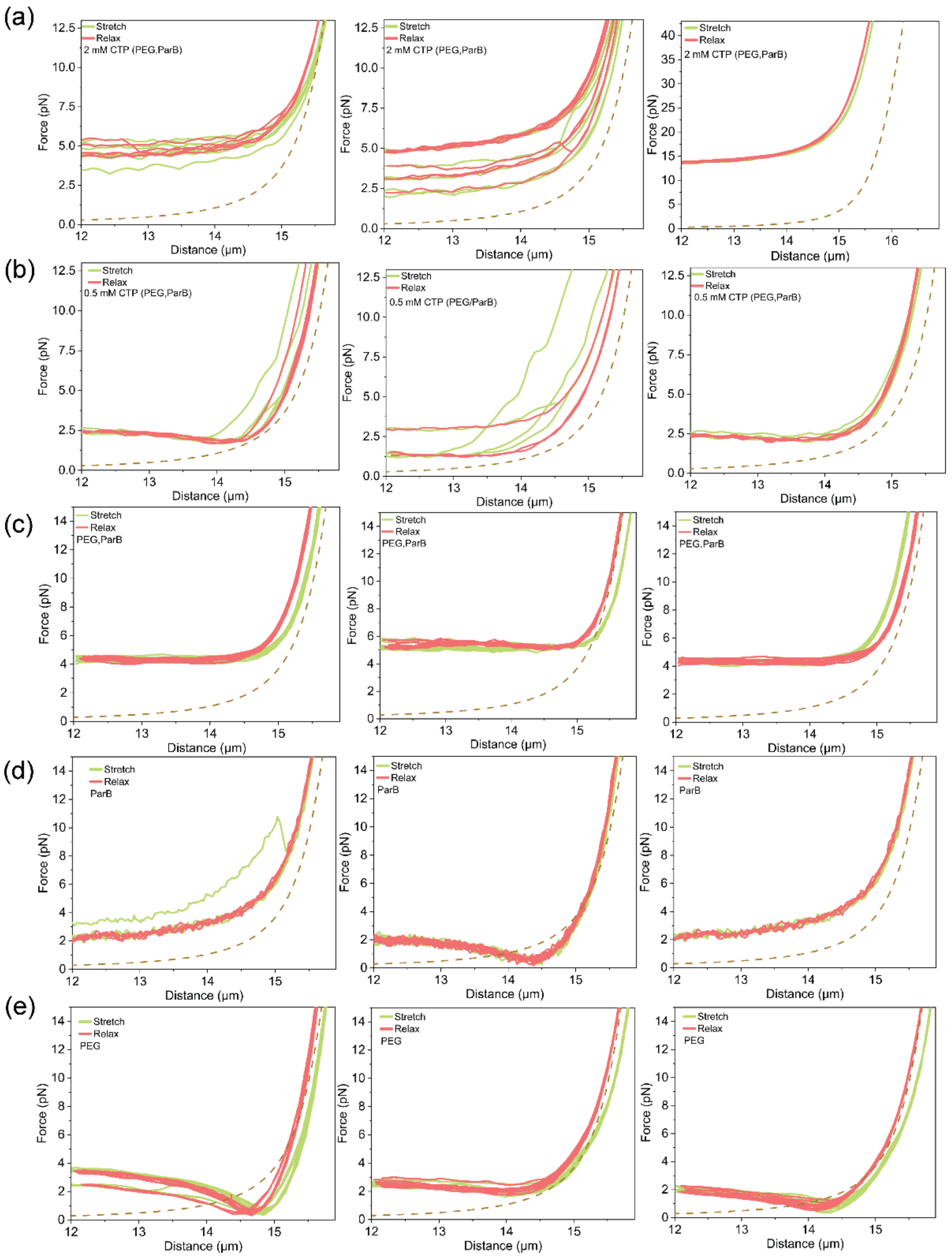


Figure S6. Force-distance curves were measured for three independent DNA tethers in different conditions (n=3 per condition).

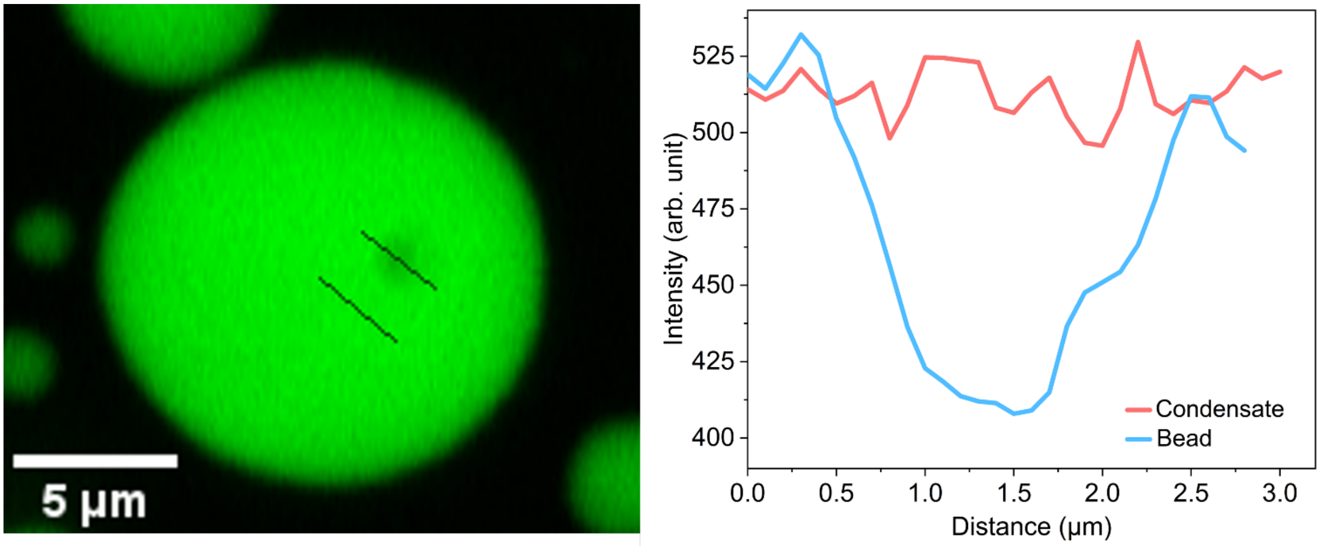


Figure S7. (Left) Fluorescence image of a condensate containing a 1 μm carboxylate bead. Two lines are marked to show intensity profiles in regions of the condensate with and without the bead. (Right) Intensity profiles across the two lines shown in the left image confirm that the beads do not adsorb ParB molecules.

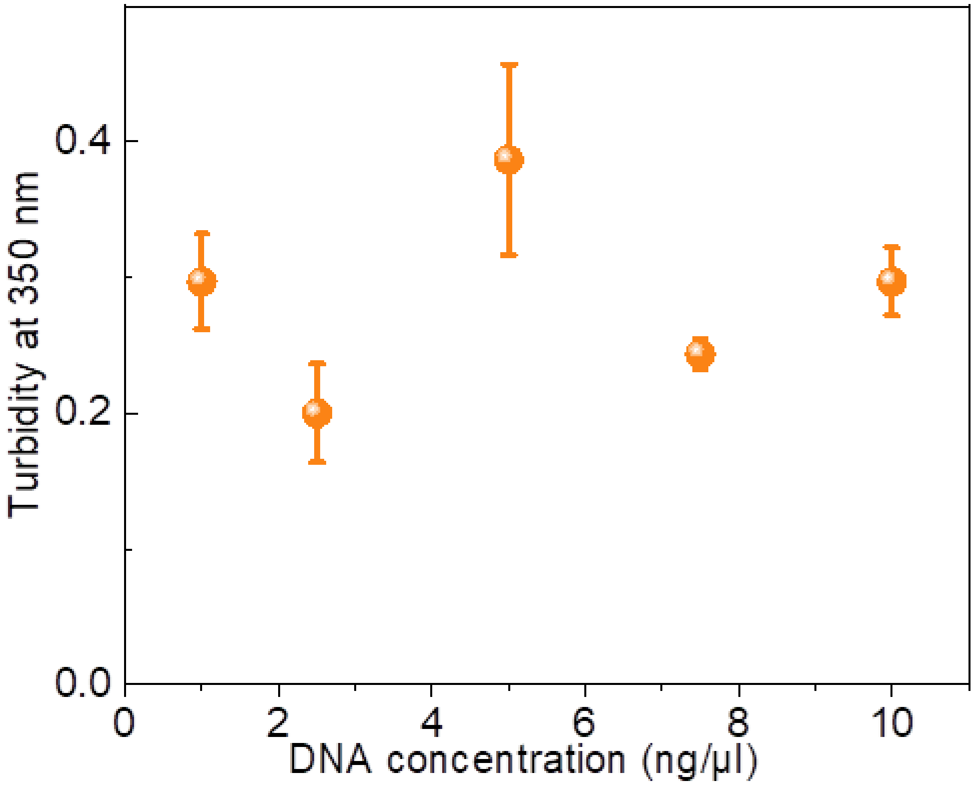


Figure S8. Turbidity of 20 µM ParB as a function of non-specific DNA in buffer with 4 mM $MgCl^{2+}$, 50 mM MPG; 2 mM CTP.

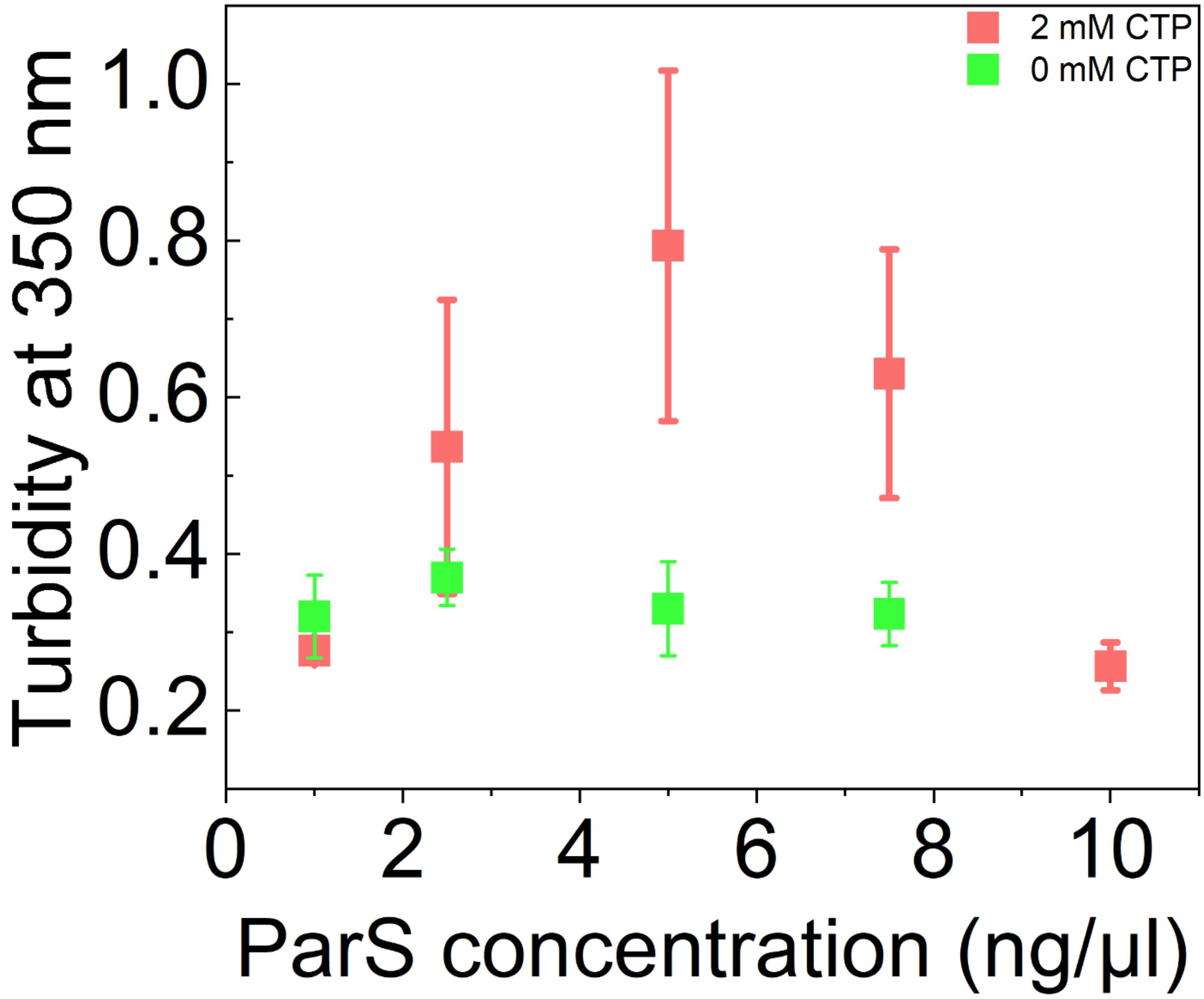


Figure S9. Turbidity of solutions containing 20 μM ParB and varying amounts of non-specific DNA in buffer with 10 mM $MgCl^{2+}$, 100 mM MPG and either 0 or 2 mM CTP.
